# QTG-Finder: a machine-learning based algorithm to prioritize causal genes of quantitative trait loci

**DOI:** 10.1101/484204

**Authors:** Fan Lin, Jue Fan, Seung Y. Rhee

**Affiliations:** Department of Plant Biology, Carnegie Institution for Science, Stanford, California 94305, USA

**Keywords:** Arabidopsis, causal gene, machine-learning algorithm, candidates, quantitative trait loci, rice

## Abstract

Linkage mapping is one of the most commonly used methods to identify genetic loci that determine a trait. However, the loci identified by linkage mapping may contain hundreds of candidate genes and require a time-consuming and labor-intensive fine mapping process to find the causal gene controlling the trait. With the availability of a rich assortment of genomic and functional genomic data, it is possible to develop a computational method to facilitate faster identification of causal genes. We developed QTG-Finder, a machine learning based algorithm to prioritize causal genes by ranking genes within a quantitative trait locus (QTL). Two predictive models were trained separately based on known causal genes in Arabidopsis and rice. With an independent validation analysis, we demonstrate the models can correctly prioritize about 65% and 60% of Arabidopsis and rice causal genes when the top 20% ranked genes were considered. The models can prioritize different types of traits though at different efficiency. We also identified several important features of causal genes including paralog copy number, being a transporter, being a transcription factor, and containing SNPs that cause premature stop codon. This work lays the foundation for systematically understanding characteristics of causal genes and establishes a pipeline to predict causal genes based on public data.

**One sentence summary:** We systematically analyzed the genomic characteristics of causal genes in QTLs and developed a novel computational tool to prioritize causal genes.

## Introduction

As the world’s population expands, food security faces a major challenge in the near future. By 2050, world population is projected to grow by 34%, which will require a 70% increase of global food production to meet the demand (FAO 2009). To catch up with the growing global food demand, it is important to improve the efficiency of arable land usage by developing better crops.

Many agriculturally and medically important traits are quantitative and controlled by multiple genetic loci. Examples include plant height, grain yield, and flowering time in plants and common disorders such as cancer, diabetes, and hypertension in humans. The variation in quantitative traits allows organisms to adapt to various environments (Baxter *et al.* 2010; Leinonen *et al.* 2013). Quantitative traits are determined by a combination of genetic complexity and environmental factors (Mackay 2001). The genetic complexity of quantitative traits comes from the involvement of multiple quantitative trait loci (QTL) and the non-additive interactions among them (Carlborg and Haley 2004; Mackay 2014). To better understand the evolutionary forces and molecular mechanisms that shape the genetic architectures of adaptive traits, we need to identify all the causal genes that contribute to most of the phenotypic variation of the traits and elucidate the molecular mechanisms of their actions. Achieving this goal will facilitate rational engineering of plant traits and more accurate prediction of the effects of the modifications on the engineered plant.

QTL linkage mapping and genome wide association study (GWAS) are two common approaches used to identify QTLs, each with its own strengths and limitations. Both mapping approaches are based on the co-segregation of a trait and genetic variants in a population. The population for linkage mapping is usually the progeny of parental plants that differ in a trait, such as an F2 population or recombinant inbred lines (Bergelson and Roux 2010). GWAS mapping uses a natural population that has a heritable variation of a trait. Compared to GWAS, linkage mapping does not suffer from issues like rare alleles and population structure (Bergelson, 2010). For example, the most significant seed dormancy QTL DOG1 identified by linkage mapping was not identified by GWAS, likely due to the rarity of the strong allele in the GWAS population (Bentsink *et al.* 2010; He 2014). Confounding population structure can cause a high false positive rate in GWAS, though some methods have been developed to ameliorate it (Price *et al.* 2010). However, efforts to correct it could result in a higher false negative rate (Brachi *et al.* 2010). Linkage mapping does not suffer from these issues, but it has a relatively lower mapping resolution and cannot identify QTLs of minor effects when the sample size is small (Martin and Orgogozo 2013; Otto and Jones 2000; Wellenreuther and Hansson 2016; Xu 2003).

For QTLs identified by linkage mapping, finding causal genes underlying them is still a big bottleneck (Bergelson and Roux 2010). In a typical rice linkage mapping, the size of a QTL can range from 200kb-3Mb, which can harbor tens to hundreds of genes depending on the mapping population and gene density (Bargsten *et al.* 2014; Daware *et al.* 2017). Even in the post-genomic era where all the genes in the genome are uncovered, identifying QTL causal genes is not straightforward since many QTLs either contain no obvious candidate genes or too many genes relevant for the trait (Nuzhdin *et al.* 1999). Therefore, despite the many QTLs that have been reported in plants, only a few have been studied at the molecular level.

Conventional fine mapping is a reliable but time-consuming and labor-intensive approach to narrow down the range of candidate genes in a QTL region. The basis of fine mapping is to create a population that has more recombination events within a QTL in order to identify a smaller genomic segment that co-segregates with the trait. However, the enormous time and labor required for creating and screening a population of progenies limits the usage of this method (Tuinstra *et al.* 1997). Depending on the frequency of recombination, thousands of progenies may need to be genotyped to get to a gene-scale resolution (Dinka *et al.* 2007). For example, 1,160 progenies were screened to identify the *Pi36* gene in rice and as many as 18,994 progenies were screened to identify the causal gene of Bph15 in rice (Liu *et al.* 2005; Yang *et al.* 2004). The high cost associated with genotyping and phenotyping makes it challenging to apply fine mapping to all QTLs.

Alternative approaches to refine the candidate list of causal genes include meta-analysis, joint linkage-association analysis, and other computational methods including machine-learning algorithms. The first two approaches require either the availability of many QTL studies on similar traits or an additional association mapping experiment (Buckler *et al.* 2009; Motte *et al.* 2014; Yin *et al.* 2017). Computational methods including machine-learning algorithms have been developed to prioritize disease associated genes and genetic variants in human (Hormozdiari *et al.* 2015; Kircher *et al.* 2014; Perez-Iratxeta *et al.* 2002; Ritchie *et al.* 2014). To distinguish disease-associated from non-associated variants, a variety of information has been used, including the effect of polymorphism (Gelfman *et al.* 2017; Kircher *et al.* 2014; Ng and Henikoff 2003), sequence conservation (Huang *et al.* 2017; Pollard *et al.* 2010), regulatory information (Deo *et al.* 2014), expression profile (Deo *et al.* 2014; Mordelet and Vert 2011), Gene Ontology (GO) (Mordelet and Vert 2011), KEGG pathway (Mordelet and Vert 2011), and publications (Perez-Iratxeta *et al.* 2002). In contrast, only two causal gene prioritization approaches are available for plants. One method was developed for GWAS in maize based on co-expression networks (Schaefer *et al.* 2018). Another method was developed for linkage mapping based on biological process GOs (Bargsten *et al.* 2014). To date, no machine-learning approaches using multiple data types have been developed to address this problem.

Here, we built a supervised learning algorithm to prioritize QTL causal genes using known causal genes in *Arabidopsis thaliana* (Arabidopsis) and *Oryza sativa* (rice) and a suite of publicly available genetic and genomic data. For each species, we trained a predictive model using features based on polymorphism data, function annotation, co-function network, and paralog copy number. By testing the models on an independent set of known causal genes, we demonstrated its efficiency in prioritizing causal genes.

## Materials and methods

### Data sources and features used in QTG-Finder

Twenty-eight features were extracted from published genome-scale data, which included polymorphism features, functional annotation features and other genomic and functional genomic features.

Arabidopsis polymorphism data of 1,135 accessions was downloaded from 1001 Genomes Project (https://1001genomes.org) (Consortium 2016) and rice polymorphism data of 3,010 cultivars was downloaded from Rice SNP-Seek Database (http://snp-seek.irri.org) (Mansueto *et al.* 2017). We used SIFT4G (v 2.4) (Ng and Henikoff 2003) and SnpEff (v 4.3r) (Cingolani *et al.* 2012) to annotate the raw polymorphism data. The number of non-synonymous SNP as annotated by SIFT4G was normalized to protein length and used as a numeric feature (normalized_nonsyn_SNP). Non-synonymous SNPs at conserved protein sequences were predicted to cause deleterious amino acid changes by SIFT4G. The presence of deleterious non-synonymous SNPs in a gene was used as a binary feature (is_nonsyn_deleterious). If a gene contained any deleterious non-synonymous SNPs, the “is_nonsyn_deleterious” feature was set to 1, otherwise it was set to 0. Other binary polymorphism features such as “is_start_lost” (start codon lost) and “is_start_gained” (start codon gained) were extracted from SnpEff annotations in the same way. For “is_SNP_cis”, the Position Weight Matrices of cis-elements were downloaded from CIS-BP database (Build 1.02) (Weirauch *et al.* 2014) and mapped to 1kb upstream of all genes in the genome using FIMO (v 4.12.0) (Grant *et al.* 2011). The cis-elements with a matching score above 55 were imported into SnpEff library to annotate the SNPs. This matching score cutoff was determined by a cross-validation as described later.

Functional annotation features were binary features based on GO (Gotz *et al.* 2008; Jones *et al.* 2014) and Plant Metabolic Network (PMN) (Schlapfer *et al.* 2017). Arabidopsis and rice genes were annotated by Blast2GO (BLAST+ 2.2.29) and InterProScan (v 5.3-46.0). The molecular function GOs were then converted to high-level functional groups such as transcription factor, receptor, kinase, transporter, and enzyme to mitigate the effect of some inaccurate annotations (Jones *et al.* 2007). Genes annotated as enzymes were further classified into 13 PMN metabolic domains such as carbohydrate metabolism and nucleotide metabolism (Schlapfer *et al.* 2017). Unclassified genes in PMN were classified as “is_other_metabolism”. Genes annotated as enzymes by GO but not present in PMN databases are enzymes involved in macromolecule metabolic process or enzymes that don’t have a specific function assigned. Since a majority of them is involved in macromolecule metabolic process, we named this group as “is_macromolecule_metabolism”.

Co-functional networks of Arabidopsis and rice were retrieved from AraNet and RiceNet (Lee *et al.* 2010; Lee *et al.* 2011). The sum of all the edge weights of a gene was used as the “network_weight” feature.

Paralog copy number (paralog_copy_number) and essential gene prediction (is_essential_gene) were taken from a previous publication (Lloyd *et al.* 2015).

### Arabidopsis and rice causal genes used for training and independent validation

For model training and cross-validation, curated causal genes from Martin and Orgogozo were used as positives for algorithm training (Martin and Orgogozo 2013). In total, 60 Arabidopsis and 45 rice causal genes were used as the initial training set. For literature validation, we performed a further literature curation and found eleven Arabidopsis and ten rice causal genes, which were not included in the Martin and Orgogozo list (Supplementary Methods).

### Algorithm training and parameter optimization

The QTG-Finder algorithm was developed in Python (v 3.6) with the ‘sklearn’ package (v 0.19.0) (Pedregosa *et al.* 2011). We developed an extended 5-fold cross-validation framework (Fig. 1a) to evaluate training performance and optimize model parameters.

**Fig. 1.**
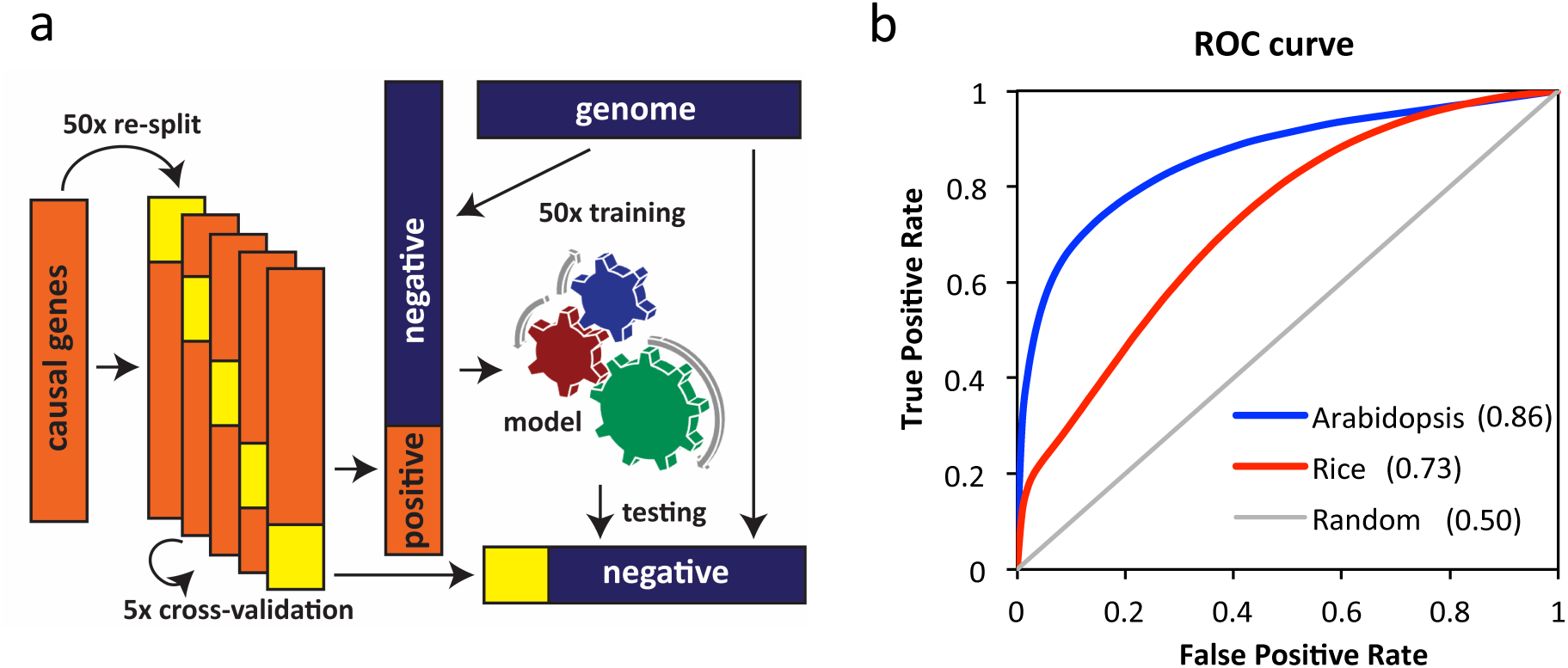
Model training and optimization based on cross-validation. (a) model training and cross-validation framework. We randomly selected negatives from the genome and iterated to maximize the combinations of training and testing data. (b) The ROC curve of Arabidopsis and rice models after parameter optimization. True and false positive rates were based on the average of all iterations. The grey diagonal line indicates the expected performance based on random guessing. The number in parentheses indicates Area Under the ROC Curve (AUC-ROC)

For the 5-fold cross validation, curated causal genes were used as positives and the other genes from the genome were used as negatives. The positives were randomly split into training and testing positives in a 4:1 ratio. Training and testing positives were combined with different sets of negative genes that were randomly selected from the rest of the genome. To increase the combination of positives and negatives, we re-split the positives 50 times randomly and selected negatives 50 times. This number of iterations ensured greater than 99% probability that every positive sample co-occurred with every negative at least once in the training or testing set during the cross-validation process. The probability of co-occurrence was calculated as Equation 1. Pco is the probability of co-occurrence of a positive and a negative in a testing or training set. N is the total number of negative samples. n is the number of negative samples selected as testing or training samples. R is the number of iterations used to re-split the positive set. C is the number of cross-validation folds that contains a positive sample. C was set to 4 for the training set and set to 1 for the testing test. S is the number of iterations to randomly select the negative set.

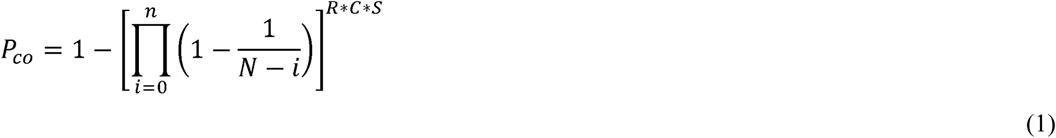

We tested different classifiers and parameters and optimized the model based on Area Under the Curve of the Receiver Operating Characteristic (AUC-ROC). The average AUC-ROC from all iterations was used to evaluate training performance. The three classifiers we tested were Random Forest, naïve Bayes, and Support Vector Machines (Cortes and Vapnik 1995; Tin Kam 1998; Zhang 2004)(Supplementary Fig. S1). For Random Forest, we tuned the number of trees and the maximum number of features for each tree based on AUC-ROC (Supplementary Fig. S2). We used 100 trees and a max_feature of 9 for Random Forest. For Support Vector Machines, RBF kernel was used and the C parameter was tuned. Random Forest was chosen for further analysis since its performance was slightly better than the other two classifiers. The ratio of positives and negatives in training data was also tuned to maximize cross-validation AUC-ROC (Supplementary Fig. S3). The best performing positives:negatives ratio was 1:20 for Arabidopsis and 1:5 for rice. For testing, a positives:negatives ratio of 1:200 was used since it is close to the average ratio of causal and non-causal genes in real QTLs.

The source code for cross-validation and any other analyses below are available at https://github.com/carnegie/QTG_Finder

### Feature importance analysis

We implemented a leave-one-out analysis to evaluate feature importance. This method was based on the change of AUC-ROC (ΔAUC-ROC) when leaving out one feature from the models. The same cross-validation framework was used for this analysis. For each iteration, we calculated AUC-ROC on the original and the leave-one-out models developed with the same training and testing datasets. The ΔAUC-ROC was calculated by subtracting the leave-one-out AUC-ROC from the original AUC-ROC. With the results from all iterations, we calculated the average ΔAUC-ROC for each feature.

### Independent literature validation

For validation, we applied the models to an independent set of causal genes that were curated from recent literature and not used for cross-validation. The models were trained by known causal genes from the initial list and negatives were randomly selected from the rest of the genome. Model training was repeated 5,000 times by resampling training negatives from the genome. With 5,000 iterations, there was >99% probability that each gene in the genome was selected at least once based on simulation. We applied the models to each of the independent causal gene and all other genes located within the QTL. All genes within the QTL were ranked based on the frequency of being predicted as a causal gene.

We calculated the probability of correctly prioritizing at least K causal genes when applying the models to a total of N QTLs with Equation 2. *p* is the probability to correctly prioritize a causal gene of a single QTL at a certain threshold. *x* is the number of causal genes being correctly prioritized.

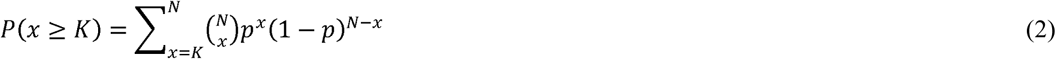

### Trait category analysis

The trait category analysis was performed in a similar way as the independent literature validation except using different training and testing sets. Each curated causal gene was tested once. For each round, one curated causal gene was removed from the training set. Then the model was trained and applied to rank the known causal gene and 200 flanking genes.

## Results

### QTG-Finder: a machine-learning algorithm to prioritize causal genes

We developed the QTG-Finder algorithm to find causal genes from QTL data and generated two predictive models in Arabidopsis and rice with the algorithm. These two species were selected for model training since they have the largest number of QTL causal genes (QTGs) that have been discovered by fine mapping and map-based cloning in plants (Martin and Orgogozo 2013). For model training, we selected 60 Arabidopsis and 45 rice causal genes as a positive set (Martin and Orgogozo, 2013, Supplementary Tables S1 and S2). The negative set was a subset of genes randomly selected from the rest of the genome. To train the models, we used 28 Arabidopsis features and 27 rice features, including polymorphisms, functional categories of genes, function interference from co-function networks, gene essentiality, and paralog copy number (Supplementary Tables S3, S4 and S5). These features were generally independent from each other (most have a Pearson’s correlation coefficient <0.2) (Supplementary Fig. S4).

We optimized the models with an extended cross-validation framework (Fig. 1a). In addition to a typical 5-fold cross-validation (Kuhn and Johnson 2013), iterations were applied to randomly select genes from the negative set and re-split the positive set in order to maximize the combinations of positives and negatives in the training and testing sets (See method).

With this framework, we evaluated the training performance with Area Under the Curve of Receiver Operating Characteristic (AUC-ROC) and optimized parameters. To find the optimal parameters, we compared the AUC-ROC of different machine-learning classifiers, modeling parameters, and the ratio of positive:negative genes in the training set (Supplementary Fig. S2, S3, and S4). Random Forest was selected as the classifier since it was less prone to over-fitting and performed better than the other classifiers tested (Supplementary Fig. S1). After optimization, AUC-ROC for the Arabidopsis and rice models were 0.86 and 0.73, respectively (Fig. 1b).

Since the positive training set used was relatively small, we also evaluated the relationship between training performance and size of the training set. The AUC-ROC increased as a larger training set was used. Interestingly, maximum gain in the AUC-ROC was achieved with 20 causal genes (Supplementary Fig. S5).

### Important features for predicting causal genes

With the optimized models, we wanted to know which features were important for causal gene prediction. Since Random Forest uses features and their interactions for classification (Touw *et al.* 2013), the importance of a feature cannot be measured by simple enrichment or depletion of a single feature in causal genes. Therefore, we evaluated feature importance based on the change of ROC-AUC (ΔROC-AUC) when excluding a feature from the model (Lloyd *et al.* 2015). When an important feature is excluded from the model, the ROC-AUC should decrease.

Here, we highlighted the six most important features out of a total of 28 features. The six most important features for Arabidopsis were paralog copy number, transporter, the number of non-synonymous SNPs normalized to protein length (normalized_nonsyn_SNP), receptor, transcription factor, and SNPs causing premature stop codon (is_stop_gained) (Fig. 2a). The six most important features for rice were paralog copy number, macromolecule metabolism, network weight sum, transcription factor, transporter, and SNPs causing premature stop codon. Four out of the six most important features were consistent between Arabidopsis and rice models, which were paralog copy number, transporter, transcription factor, and SNPs causing premature stop codon.

**Fig. 2.**
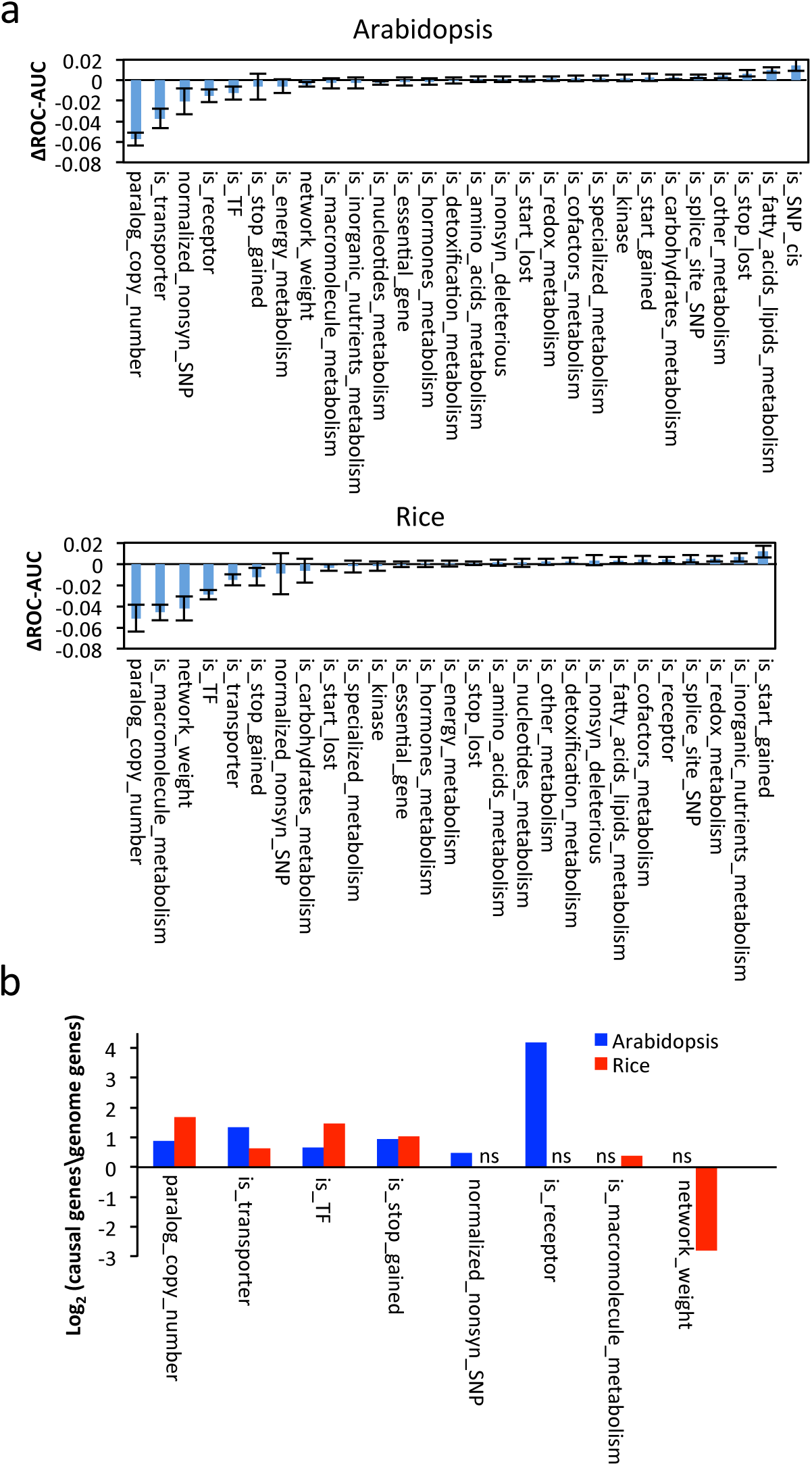
Important features of causal genes and their enrichment or depletion relative to the genome background (a) Feature importance as indicated by the change of AUC-ROC (ΔAUC-ROC) when excluding each feature. The ΔAUC-ROC indicates the average value of all iterations. Error bars indicate standard deviation. The features with a name that starts with “is_” are binary variables. (b) The enrichment or depletion of the top 6 features in Arabidopsis and rice models. The enrichment/depletion were indicated by the ratio of causal genes to genome background. ns, not shown because the feature is not one of the top 6 features in that species

For the six most important features in Arabidopsis and rice, we examined their ratio in known causal genes versus randomly selected genes in the genome (Fig. 2b). Compared to other genes in the genome, the causal genes tended to have more paralogs, higher frequency of being a transporter or a transcription factor, and higher frequency of containing SNPs that cause premature stop codons in both species.

The rest of the features contributed less to, but did not impair, model performance to a large degree (ΔROC-AUC< 0.02). Since there was no strong evidence that they impair prediction, we did not remove them from the models for further analysis.

### Validating QTG-Finder by ranking an independent set of QTL genes

To assess the predictability of QTG-Finder models, we searched the literature for a separate set of known causal genes from the initial training set. We found eleven Arabidopsis and ten rice genes that are likely causal genes underlying QTLs when interpreting linkage mapping with additional evidence such as functional complementation, fine mapping, joint linkage-association analysis or genetic analyses (Supplementary Table S6). These causal genes were not used for model training or cross-validation.

To examine model performance, we applied the QTG-Finder models to this new set of causal genes. For each known causal gene, we ranked all genes in the QTL region based on the frequency of being predicted as a causal gene from 5,000 iterations. Since the number of genes in a QTL region varies, we used a gene’s rank percentile for evaluation. The rank percentile of a gene indicates the percentage of QTL genes that had higher ranks than the gene of interest.

Based on the rank of these known causal genes, we evaluated model performance at different cutoffs. We calculated the percentage of known causal genes included in the top 5%, 10%, and 20% of the prioritized genes within a QTL (Fig. 3a). The top 20% of the ranked genes included seven Arabidopsis (∼64%) and six rice (∼60%) causal genes. With a more stringent cutoff of 5%, four Arabidopsis (∼27%) and three rice (∼30%) causal genes were prioritized.

**Fig. 3.**
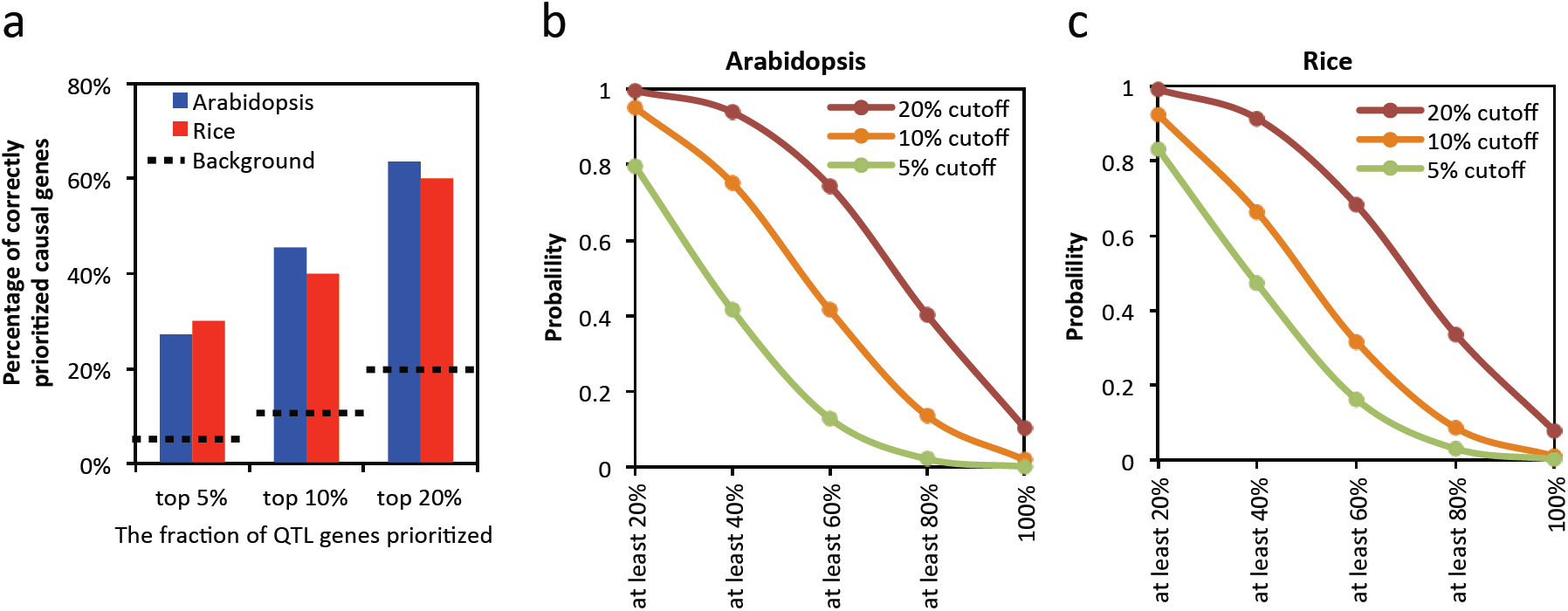
Model performance at different thresholds (a) Percentage of correctly prioritized causal genes of a single QTL at different rank thresholds. Dashed lines indicate the background of random selections. (b-c) The probability of correctly prioritizing at least X% of causal genes when analyzing multiple QTLs simultaneously

Most linkage mapping studies identify multiple QTLs. We therefore calculated a theoretical model performance on identifying causal genes from multiple QTLs simultaneously, which we defined as the probability of identifying at least X% of all causal genes when applying the model to all QTLs of a trait (Fig. 3b and c). For example, assuming there were five QTLs of a trait identified by a linkage mapping study and each QTL contained one causal gene. For the Arabidopsis model, the probability of identifying at least one causal gene would be 99% when the top 20% genes of all QTLs were tested experimentally. The probability of identifying all five causal genes would be 10% when the top 20% cutoff was used. We further compared the performance of all three cutoffs, top 20%, top 10%, and top 5%. The probability of identifying at least one out of five causal genes would be no less than 80% for all three cutoffs. The probability to correctly prioritize at least four out of five causal genes would be 40% (for top 20%), 14% (for top 10%), and 2% (for top 5%). Therefore, a less stringent cutoff (top 20%) performs much better than a more stringent cutoff if one is interested in finding most of the causal genes or causal genes of a particular QTL. However, if the goal is to identify any causal gene, then screening the top 5% of all QTLs may be a more strategic approach since fewer candidate genes need to be tested experimentally.

### Trait type preference of QTG-Finder models

Since the training set included genes for different types of traits at an imbalanced ratio, we wanted to know how QTG-Finder models would work for each type of traits (Fig. 4a). The independent validation described above was based on causal genes related to plant development and disease resistance (Supplementary Table S6). However, this validation set was not large enough for a systematic analysis and did not have any abiotic-stress-related causal genes. Therefore, we performed a rank analysis for different trait categories using the known causal genes from the initial training set (60 for Arabidopsis and 45 for rice). For this rank analysis, each causal gene was taken out from the training set once and used for a rank test. The single causal gene and its 200 neighboring genes in the genome were used as a testing set. We applied the models to each testing set to obtain the rank for each causal gene. Then we calculated the average rank for the causal genes in the four trait categories: development, abiotic stress, biotic stress and “other”. The “other” category included traits in seed hull color, oil composition, necrosis, etc.

**Fig. 4.**
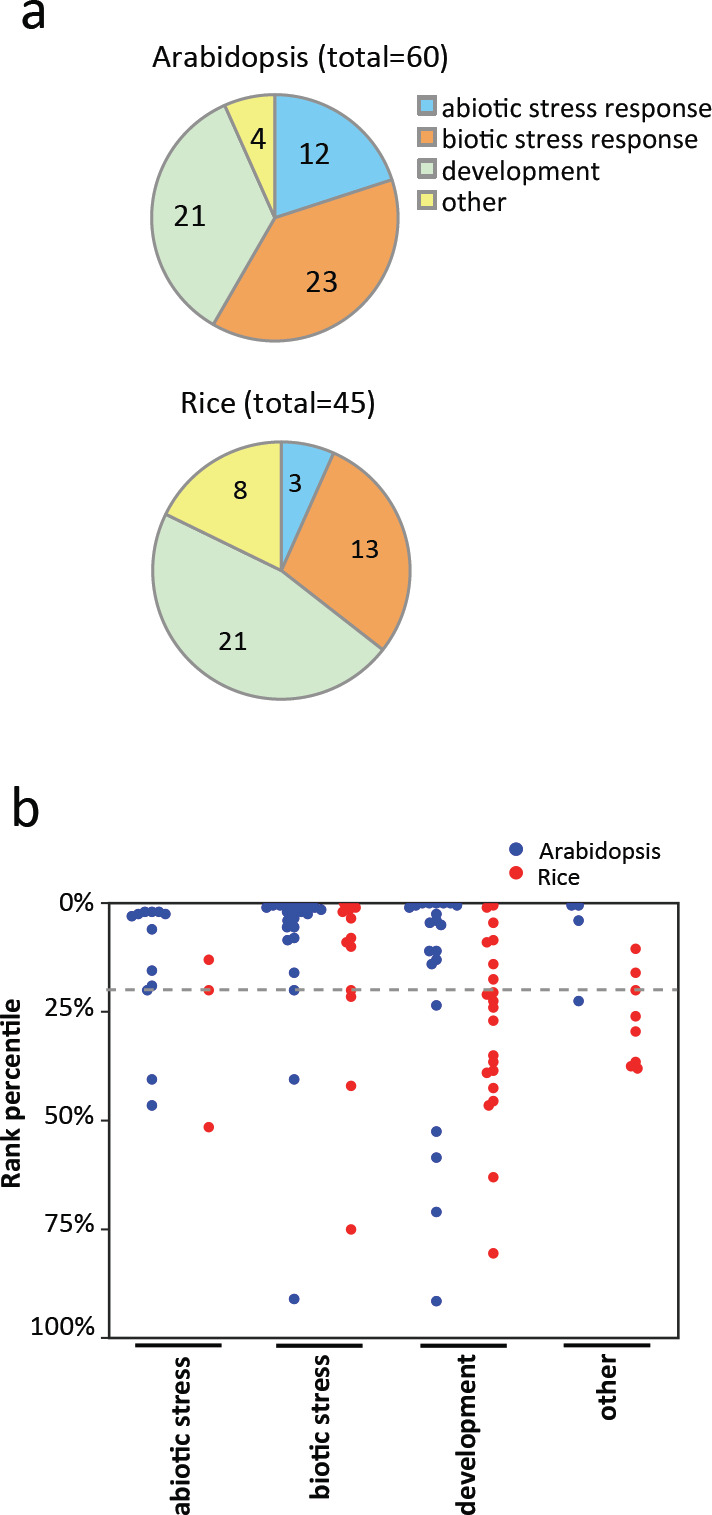
(a) Trait categories of known causal genes from the training set. (b) The rank percentile of causal genes of different trait categories. Each causal gene and 200 neighboring genes were used as testing set once. All other known causal genes were used as training set. Each dot indicates a known causal gene. The grey dashed line indicates 20% rank percentile. The trait categories of causal genes are defined in Tables S1 and S2

Performance of the models was not the same for different trait categories. Both abiotic and biotic stress traits had better performance than developmental traits (Fig. 4b). In addition, the Arabidopsis model performed slightly better than the rice model for all trait categories. This trait category analysis can guide users to determine rank cutoffs when applying models to different types of traits.

## Discussion

Linkage mapping is a useful tool to identify the genomic regions responsible for many agriculturally and medically important traits. However, it is not straightforward to identify the genes that cause the trait variation from these genome regions. The discovery of causal genes still requires time-consuming and labor-intensive fine mapping. In this study, we developed a machine-learning algorithm to reduce the number of candidates to be tested experimentally in order to accelerate the discovery of causal genes.

### A machine-learning algorithm to prioritize QTL causal genes

Several causal variant or gene prioritization methods have been developed for human data but not many in plants (Bargsten *et al.* 2014; Jagadeesh *et al.* 2016; Kircher *et al.* 2014; Schaefer *et al.* 2018). Most prioritization methods have been developed for GWAS mapping in human, an organism where linkage mapping cannot be performed. However, linkage mapping can capture rare alleles and has been broadly used to study quantitative traits of livestock, crops, and model organisms. A causal gene prioritization is especially helpful for large QTLs identified by linkage mapping, which can constitute tens to hundreds of genes. One method has been developed in rice to prioritize causal genes for linkage mapping (Bargsten *et al.* 2014). This method is based on the hypothesis that causal genes from multiple QTLs of the same trait are more likely to have the same biological process GO terms, and therefore genes with overrepresented biological process GOs were prioritized as causal genes. However, this method gives no predictions for ∼15% of traits and lack an unbiased performance evaluation since the same set of causal genes was used to determine cutoff and evaluate performance.

In this study, we built a supervised learning algorithm using multiple features and validated its efficacy with an independent dataset from the literature. The models could accelerate the discovery of causal genes by ranking all the genes in a QTL region and prioritizing the top 5%, 10%, or 20% genes, which are most likely to contain the causal gene, for experimental testing. Based on an assessment using independent data in the literature, we calculated the performance when applying the models to all QTLs of a trait and compared three cutoffs (top 5%, 10%, and 20%). The less stringent cutoff (top 20%) had a higher chance to find more causal genes (Fig. 3b and c) but yielded more candidates that needed to be tested by experiments. The more stringent cutoff (top 5%) had a lower chance to find all causal genes but yielded a smaller set of candidates to test. The probability for the models to find at least one causal gene is high for all three cutoffs. If the goal were to find one or more causal genes for functional studies and the particular QTL regions did not matter, the 5% cutoff would be more efficient. If the goal were to discover all causal genes and understand the genetic architecture of a trait, the 20% cutoff would be better. Similarly, if a particular QTL were of interest for discovering the underlying causal gene, the 20% cutoff would be better.

There are several conceptual and practical advantages of QTG-Finder algorithm. First, this algorithm combines multiple types of publically available data including polymorphisms, function annotations, co-function network and other genomic data, which have not been applied to prioritize causal genes from linkage mapping studies. Second, models were trained on causal genes from various traits and can be applied to several types of traditional traits, though the prioritization efficiency was not equivalent. Third, validation from the literature provides guidance on what proportion of genes to prioritize in practice rather than arbitrarily selecting a threshold. Fourth, the models treat each QTL independently and have the flexibility to prioritize a specific QTL of interest.

Two limitations of this study are the small number of known causal genes in plants and the impurity of negative set used for model training. We used 60 Arabidopsis and 45 rice causal genes that have been verified by map-based-cloning as a positive dataset. Even though they are of high quality, this positive dataset may not be large enough to represent all the features of causal genes. There could still be other important features of causal genes that we were not able to capture with this small dataset. The negative set was composed of genes randomly selected from the rest of the genome. Though we excluded known causal genes, there could still be some uncharacterized causal genes. As a result of these limitations, 20% cutoff will still yield ∼100 candidates for large QTLs, which is challenging for genetic characterization unless at least a medium-throughput phenotyping method is available. Fortunately, plant science is entering an era of high-throughput phenotyping with advances in automation, computation and sensor technology (Araus *et al.* 2018; Fahlgren *et al.* 2015). Our study establishes an extendable framework that can be easily updated with new training datasets and features. As more causal genes are uncovered, the new data can be easily incorporated to improve the models.

### Important features for predicting QTL causal genes

Many causal genes were repeatedly found to cause phenotypic variation of similar traits, which is also known as genetic hotspots of phenotypic variation or gene reuse (Martin and Orgogozo 2013). By examining 1,008 causative alleles in animals, plants, and yeasts, Martin and Orgogozo found *de novo* mutations to occur repeatedly at certain genes or orthologous loci and causing similar phenotypic variations either among lineages or within a single lineage. The prevalence of gene reuse suggests that causal genes are likely to have some genetic and genomic characteristics that allow them to be repeatedly used for phenotypic variation. The mechanism for gene reuse is not clear but it may be influenced by factors such as the availability of standing genetic variation, mutation rate, pleiotropic constraint, and epistatic interactions of a gene (Conte *et al.* 2015; Conte *et al.* 2012).

While many QTL causal genes have been cloned, their features have not been systematically examined before. Instead of evaluating each feature individually, we trained Random Forest models and evaluated feature importance for all features by adopting the leave-one-out strategy. Several of the most important features were consistent between Arabidopsis and rice models: containing SNPs that cause a premature stop codon, paralog copy number, being a transporter, and being a transcription factor.

We extracted polymorphism features from re-sequencing data, which provide more information about the existence of standing genetic variation in the species than the polymorphism data used for linkage mapping, which typically comes from two parental lines. DNA polymorphisms such as nonsense SNPs, deleterious non-synonymous SNPs, SNPs at cis-regulatory elements, and SNPs at splice junctions have been used as features to classify causal and non-causal variants of human diseases (Jagadeesh *et al.* 2016; Kircher *et al.* 2014). These SNPs can directly affect the function or expression of a gene and therefore are more likely to be causal than the rest of the SNPs. Our results indicate Arabidopsis and rice causal genes were more likely to carry a SNP that causes premature stop codon (nonsense SNP) than an average gene in the genome. We also found Arabidopsis causal genes were more likely to have more non-synonymous SNPs than an average gene in the genome. Besides the high impact SNPs in coding regions, we also examined polymorphisms in non-coding regions since about 90% of human trait/disease-associated SNPs are located outside of coding regions (Hindorff *et al.* 2009). The SNPs at cis-regulatory elements did not show a high feature importance in our algorithm, although this might be due to limited exploration of non-coding sequences in plants. The CIS-BP database contains cis-elements of 44% of the transcription factors in Arabidopsis (Weirauch *et al.* 2014). Developing a more accurate and complete map of functional non-coding regions based on conserved noncoding sequences (Van de Velde *et al.* 2014) will potentially make non-coding polymorphism features more useful for prioritizing causal genes.

Paralogs contribute to the evolution of plant traits by providing functional divergence that gives plants the potential to adapt to complex environments (Panchy *et al.* 2016). Through evolution, genes involved in signal transduction and stress response have retained more paralogs while essential genes like DNA gyrase A have retained fewer paralogs (Lloyd *et al.* 2015; Panchy *et al.* 2016). By acquiring new functions or sub-functions, paralogs allow plants to sense and handle different environmental conditions in a more comprehensive and adjustable way. For example, the eight paralogous heavy metal ATPases (HMAs) in Arabidopsis are all involved in heavy metal transport but have different substrate preference, tissue expression patterns, and subcellular compartment locations (Takahashi *et al.* 2012). Three of them, HMA3, HMA4, HMA5, are known causal genes of QTLs identified by linkage mapping. The known causal genes we analyzed have more paralog copies than other genes in the genome. This may suggest that many plant causal genes are playing a role in providing more phenotypic tuning parameters to allow plants to adapt to complex environments.

Plant transporters are involved in nutrient uptake, response to abiotic stresses, pathogen resistance, and other plant-environment interactions (Conde *et al.* 2011; Doidy *et al.* 2012). Polymorphisms in transporters play an important role in local adaptation since many transporters are directly involved in environment responses (Baxter *et al.* 2010; Turner *et al.* 2010). For example, in *Arabidopsis lyrata*, the polymorphisms most strongly associated with soil type are enriched in metal transporters (Turner *et al.* 2010). We observed a higher frequency of causal genes being transporters than the average gene in the genome. Causal transporters that contribute to trait variation may have a more important role in local adaptation than other transporters.

Transcription factors were enriched in causal genes not only in plants but also in other organisms (Martin and Orgogozo 2013). This enrichment may be due to an ascertainment bias since linkage mapping tends to identify genes with large effects (Martin and Orgogozo 2013). Since QTG-Finder focuses on prioritizing the causal genes identified by linkage mapping, this feature is useful in distinguishing them from other causal genes such as the medium-effect genes that can be detected by GWAS but not by linkage mapping.

Overall, QTG-Finder is a novel machine-learning pipeline to prioritize causal genes for QTLs identified by linkage mapping. We trained QTG-Finder models for Arabidopsis and rice based on known causal genes from each species, respectively. By utilizing information like polymorphisms, function annotations, co-function networks, and paralog copy numbers, the models can rank QTL genes to prioritize causal genes. Our independent literature validation demonstrates that the models can correctly prioritize about 65% of causal genes for Arabidopsis and 60% for rice when the top 20% of ranked QTL genes were considered. The algorithm is applicable to any traditional quantitative traits but the performance was different for each trait type. Since QTG-Finder is a machine-learning based pipeline, extending the training set and adding features can easily expand and improve the models. We envision that frameworks like QTG-Finder can accelerate the discovery of novel quantitative trait genes by reducing the number of candidate genes and efforts of experimental testing.

## Supporting information

Supplementary_materals

## Author Contributions

Conceptualization, S.Y.R.; Methodology, J.F., F.L., and S.Y.R.; Investigation, F.L. and J.F.; Formal Analysis, F.L. and J.F.; Writing–Original Draft, F.L.; Writing–Review & Editing, F.L., S.Y.R., and J.F.; Funding Acquisition, S.Y.R.; Resources, S.Y.R.; Supervision, S.Y.R.

## Acknowledgements

We thank Dr. John Lloyd and Dr. Shin-Han Shiu for sharing the data of rice essential gene prediction. We thank Kevin Radja for testing the source code and giving useful comments.

