## Supplementary_materals for "QTG-Finder: a machine-learning based algorithm to prioritize causal genes of quantitative trait loci"

#### List of Contents

**Supplementary Methods**
**Supplementary Table S1** Curated Arabidopsis causal genes used for model training and cross-validation
**Supplementary Table S2** Curated rice causal genes used for model training and cross-validation
**Supplementary Table S3** Causal gene features and their frequency and value in causal genes and the genome background
**Supplementary Table S4** Features used for the Arabidopsis model
**Supplementary Table S5** Features used for the rice model
**Supplementary Table S6** Evaluating model performance based on the rank of an independent set of known causal genes.
**Supplementary Fig. S1** Comparing ROC curves of classifiers
**Supplementary Fig. S2** Random Forest hyperparameters
**Supplementary Fig. S3** The ratio of positives:negatives in training set
**Supplementary Fig. S4** Correlations among features
**Supplementary Fig. S5** The relationship between training performance and the number of training positives used
**References**

### Supplementary Methods

#### Curation of causal genes for training and cross-validation

We curated the causal genes (Supplementary Tables S1 and S2) based on the list of causal alleles in (Martin and Orgogozo 2013). The curated causal genes only included genes that were cloned. Alleles of the same causal genes identified from different parental lines were merged into one causal gene entry for training. Since only gene names were available for these causal genes in the original list (Martin and Orgogozo 2013), we searched the literature to find gene IDs associated with sequences. When the gene ID was not provided in the literatures, we obtained the sequence of the gene from GenBank (https://www.ncbi.nlm.nih.gov/genbank/) or UniProt (https://www.uniprot.org) and performed BLAST against the Arabidopsis Information Resource (https://www.arabidopsis.org/Blast/) or the MSU Rice Genome Annotation Project Database (http://rice.plantbiology.msu.edu/ analyses_search_blast.shtml) to get the gene IDs.

#### Curation of causal genes for independent validation

For literature validation, we performed a further literature curation and found eleven Arabidopsis and ten rice causal genes, which were not included in the Martin and Orgogozo list. It included causal genes that have been cloned or indicated by joint linkage-association analysis or indicated by genetic analyses (Azizi *et al.* 2016; Barth and Jander 2006; Bennett *et al.* 2006; Chardon *et al.* 2013; Ehrenreich *et al.* 2007; Fan *et al.* 2016; Fukuoka *et al.* 2014; Gao *et al.* 2016; Guo *et al.* 2015; Hu *et al.* 2008; Huang *et al.* 2012; Itoh *et al.* 2010; Liu *et al.* 2017; Motte *et al.* 2014; Oikawa *et al.* 2015; Qi *et al.* 2008; Qiu *et al.* 2007; Rai *et al.* 2011; Riefler *et al.* 2006; Sato *et al.* 1999; Sharma *et al.* 2005; Yuan *et al.* 2016; Zeng *et al.* 2013).

### Supplementary Tables

**Supplementary Table S1** Curated Arabidopsis causal genes used for model training and cross-validation.

| Gene Name | Gene ID | Gene Function | Trait | Trait category | Reference PMID |
|---|---|---|---|---|---|
| APR2 | AT1G62180 | enzyme | shoot sulfate content | abiotic stress response | 17589509 |
| ACD6 | AT4G14400 | membrane protein | leaf initiation and necrosis | development | 20520716, 20336072 |
| AOP3 | AT4G03050 | enzyme | glucosinolate accumulation | biotic stress response | 11251105, 19737743, 21857804 |
| AT5G41740 | AT5G41740 | R-protein with leucine-rich repeats | necrosis | other | 17803357 |
| AT5G41750 | AT5G41750 | R-protein with leucine-rich repeats | necrosis | other | 17803357 |
| AtHKT1 | AT4G10310 | transporter ion | salt tolerance | abiotic stress response | 17140289, 21085628 |
| Brevis radix (BRX) | AT1G31880 | TF | root length | development | 15031265 |
| Cryptochrome 2 (CRY2) EDI allele | AT1G04400 | chromophore protein | flowering time and other pleiotropic effects | development | 11726930, 15280248, 15248119, 14605225 |
| CYCD5;1 | AT4G37630 | cyclin-dependent kinase | cell division | development | 22392991 |
| CYP81F2 | AT5G57220 | enzyme (P450) | glucosinolate metabolism | biotic stress response | 19293369 |
| DOG1 (DELAY OF GERMINATION 1) | AT5G45830 | unknown | seed germination | development | 17065317, 22231484, 20336072 |
| EARLY FLOWERING 3(ELF3) | AT2G25930 | Circadian oscillator | flowering | development | 20838594, 21857804, 20713464, 23129635 |
| Epithiospecifier Modifier1 (ESM1) | AT3G14210 | enzyme associated | glucosinolate | biotic stress response | 16679459 |
| Epithiospecifier protein (ESP) | AT1G54040 | enzyme | glucosinolate | biotic stress response | 11752388 |
| ERECTA | AT2G26330 | receptor (RTK-LRR) | plant and leaf architecture, transpiration | development | 16007076, 20374533, 21368205 |
| FLC (Flowering | AT5G10140 | TF (MADS) | flowering | development | 22865739 |

| | | | | | |
|---|---|---|---|---|---|
| Locus C) | | | | | |
| FLM (MAF1) | AT1G77080 | TF (MADS) | flowering | development | 15695584 |
| Flowering locus T (FT) | AT1G65480 | RAF kinase inhibitor | flowering | development | 17158798 |
| FPN2 | AT5G03570 | transporter | nickel tolerance | abiotic stress response | 19861554 |
| FRD3 (FERRIC REDUCTASE DEFECTIVE3) | AT3G08040 | transporter | iron transport | abiotic stress response | 23236296 |
| Frigida (FRI) | AT4G00650 | nuclear regulatory protein | flowering time | development | 11030654, 12140238, 12805638, 15908596 |
| Frigida like 1 (FRL1) | AT5G16320 | nuclear regulatory protein, coiled-coil domain-containing protein | flowering time | development | 17056759 |
| Frigida like 2 (FRL2) | AT1G31814 | nuclear regulatory protein, coiled-coil domain-containing protein | flowering time | development | 17056759 |
| GIBBERELLIC ACID REQUIRING 1 (GA1) | AT4G02780 | enzyme | plant morphology (flowers) | development | 22510148 |
| heavy metal atpase3 (HMA3) | AT4G30120 | transporter | cadmium accumulation | abiotic stress response | 22969436 |
| heavy metal atpase4 (HMA4) | AT2G19110 | transporter | cadmium accumulation | abiotic stress response | 17434989, 18425111 |
| heavy metal atpase5 (HMA5) | AT1G63440 | transporter | copper accumulation | abiotic stress response | 18701674 |
| HUA2 | AT5G23150 | Signalling RPR domain protein, putative mRNA processing factor | flowering time, shoot morphology | development | 17764945 |
| KCS18 | AT4G34520 | enzyme | oil composition | other | 23145136 |
| MADS AFFECTING FLOWERING 2 (MAF2) | AT5G65050 | TF (MADS) | flowering | development | 20551443 |
| MAM1 | AT5G23010 | enzyme | glucosinolate | biotic stress response | 11706188, 19737743, 21857804, 23042895 |
| metal tolerance protein1 | AT2G46800 | transporter | zinc concentration | abiotic stress response | 15255871, 20419142 |

| | | | | | |
|---|---|---|---|---|---|
| Molybdenum transporter1 (MOT1) | AT2G25680 | transporter | molybdenum content | abiotic stress response | 18454190 |
| mucilage-modified 2 (mum2) | AT5G63800 | enzyme | mucilage formation | other | 18165330 |
| phytochrome A (PHYA) | AT1G09570 | chromophore protein | light sensitivity | development | 11726931 |
| phytochrome B (PHYB) | AT2G18790 | chromophore protein | light sensitivity | development | 18287016 |
| phytochrome C (PHYC) | AT5G35840 | chromophore protein | light sensitivity | development | 16732287 |
| phytochrome D (PHYD) | AT4G16250 | chromophore protein | light sensitivity, increased petiole length, reduced cotyledon area and anthocyanin accumulation in seedling stems | development | 9286109 |
| RAS1 | AT1G09950 | microProtein | salt tolerance | abiotic stress response | 20212128 |
| resistant to methyl viologen 1 (RMV1) | AT5G05630 | transporter | polyamine uptake | abiotic stress response | 22492932 |
| RLM1 | AT1G64070 | R-protein with leucine-rich repeats | pathogen resistance | biotic stress response | 16623885 |
| RLM3 | AT4G16990 | R-protein with leucine-rich repeats | pathogen resistance | biotic stress response | 18397376 |
| RPM1 | AT3G07040 | R-protein with leucine-rich repeats | pathogen resistance | biotic stress response | 16623885 |
| RPP13 | AT3G46530 | R-protein with leucine-rich repeats | pathogen resistance | biotic stress response | 15082565 |
| RPP4 | AT4G16860 | R-protein with leucine-rich repeats | pathogen resistance | biotic stress response | 11846877 |
| RPP5 | AT4G16950 | R-protein with leucine-rich repeats | pathogen resistance | biotic stress response | 9212464, 11846877, 20479233 |
| RPP8 | AT5G43470 | R-protein with leucine-rich repeats | pathogen resistance | biotic stress response | 9811794 , 20479233 |
| RPS2 | AT3G03600 | R-protein with leucine-rich repeats | pathogen resistance | biotic stress response | 9874813 |
| RPS4 | AT5G45250 | R-protein with leucine-rich repeats | pathogen resistance | biotic stress response | 11846877 |
| RPS5 | AT1G12220 | R-protein with leucine-rich repeats | pathogen resistance | biotic stress response | 9212464, 11846877, 20479233 |
| RRS1 | AT5G45260 | R-protein with leucine-rich repeats | pathogen resistance | biotic stress response | 19519800, 19686535 |
| WRR4 | AT1G56510 | R-protein with leucine-rich repeats | pathogen resistance | biotic stress response | 18624640 |
| CBF gene cluster | AT4G25490 | TF (MADS) | cold resistance | abiotic stress | 21421342 |

| | | | | | |
|---|---|---|---|---|---|
| | | | | response | |
| RAC1 | AT1G31540 | R-protein with leucine-rich repeats | pathogen resistance | biotic stress response | 15242165 |
| RPP1-WsA | AT3G44670 | R-protein with leucine-rich repeats | pathogen resistance | biotic stress response | 9811793 |
| RPP1-WsB | AT3G25510 | R-protein with leucine-rich repeats | pathogen resistance | biotic stress response | 9811793 |
| RPP1-WsC | AT3G44480 | R-protein with leucine-rich repeats | pathogen resistance | biotic stress response | 9811793 |
| RPP2A | AT4G19500 | R-protein with leucine-rich repeats | pathogen resistance | biotic stress response | 15165183 |
| RPP2B | AT4G19510 | R-protein with leucine-rich repeats | pathogen resistance | biotic stress response | 15165183 |
| GLABROUS1 | AT3G27920 | TF (MYB) | trichome (leaves) | development | 11504855, 17217357 |

**Supplementary Table S2** Curated rice causal genes used for model training and cross-validation.

| Gene name | ID | Gene function | Trait | Trait category | Reference PMID |
|---|---|---|---|---|---|
| Alk / Starch Synthase II | LOC_Os06g12450 | enzyme | grain cooking texture | other | 12579422, 16027975, 20972439 |
| BADH2 | LOC_Os08g32870 | enzyme | fragrance | other | 17129318, 19706531 |
| Ehd1 | LOC_Os10g32600 | TF | flowering time | development | 15078816 |
| EARLY FLOWERING 3/Hd17 | LOC_Os06g05060 | Circadian oscillator | flowering time | development | 22399582 |
| Ghd7 | LOC_Os07g15770 | TF (CO-like) | flowering time, plant morphology (inflorescence) | development | 18454147 |
| Hd6a | LOC_Os03g55389 | Protein kinase CK2 | flowering time | development | 11416158 |
| PROG1 | LOC_Os07g05900 | TF | plant architecture | development | 18820699, 18820696, 23034647 |
| qPE9-1 | LOC_Os09g26999 | keratin-associated protein | plant and inflorescence architecture | development | 19546322 |
| qSH1 | LOC_Os01g62920 | TF BEL1-type homeobox gene (presumptive transription factor) | seed shattering | development | 16614172 |
| se5 | LOC_Os06g40080 | chromophore protein | flowering time | development | 10849355 |
| Shattering1 - OsSh1 | LOC_Os03g44710 | TF (YABBY-like) | seed shattering | development | 22581231, 21695282, |

| | | | | | |
|---|---|---|---|---|---|
| | | | | | 22158310 |
| shattering4 | LOC_Os04g57530 | TF (homeobox) | seed shattering | development | 16527928 |
| SKC1 =OsHKT1 | LOC_Os06g48810 | transporter (HKT-type) | salt homeostasis and salt tolerance | abiotic stress response | 16155566 |
| heavy metal atpase3 (HMA3) | LOC_Os07g12900 | transporter | metal tolerance | abiotic stress response | 20823253 |
| Nramp aluminum transporter1 | LOC_Os02g03900 | transporter | metal tolerance | abiotic stress response | 21829395 |
| Pi-ta | LOC_Os12g18360 | R-protein with leucine-rich repeats | pathogen resistance | biotic stress response | 11090207, 21695282, 22158310 |
| Pi2 (Nbs4-Pi2) | LOC_Os06g17920 | R-protein with leucine-rich repeats | pathogen resistance | biotic stress response | 17073304 |
| Pi9 (= Nbs2-Pi9) | LOC_Os06g05359 | R-protein with leucine-rich repeats | pathogen resistance | biotic stress response | 16387888 |
| Piz-t | LOC_Os06g17900 | R-protein with leucine-rich repeats | pathogen resistance | biotic stress response | 17073304 |
| Pi37 | LOC_Os01g57310 | R-protein with leucine-rich repeats | pathogen resistance | biotic stress response | 17947408 |
| Pid3 | LOC_Os06g22460 | R-protein with leucine-rich repeats | pathogen resistance | biotic stress response | 21621742 |
| Xa1 | LOC_Os04g53160 | R-protein with leucine-rich repeats | pathogen resistance | biotic stress response | 9465073 |
| Xa21 | LOC_Os01g56470 | R-protein with leucine-rich repeats | pathogen resistance | biotic stress response | 8525370 |
| Xa26 | LOC_Os04g13640 | R-protein with leucine-rich repeats | pathogen resistance | biotic stress response | 14756760 |
| OsGA20ox1 | LOC_Os03g63970 | enzyme | seedling vigor | development | 22481119 |
| GRAIN INCOMPLETE FILLING 1 | LOC_Os04g33740 | enzyme | grain weight | development | 18820698 |
| GS5 | LOC_Os05g06660 | enzyme | grain size | development | 22019783 |
| GW2 | LOC_Os02g14720 | enzyme E3 ubiquitin ligase | grain size | development | 17417637 |
| OsCKX2=Gn1a | LOC_Os01g10110 | enzyme | grain yield | development | 15976269 |
| OsPPKL1/qGL3 | LOC_Os03g44500 | signal transduction | grain size | development | 23236132 |
| OsSPL14 / WFP | LOC_Os08g39890 | TF | grain yield | development | 20495565, 20495564 |
| OsSPL16 | LOC_Os08g41940 | TF | grain size and shape | development | 22729225 |
| qSW5 | LOC_Os05g09520 | unknown | grain size | development | 18604208, 20972439 |
| Sd1 (=GA20ox-2) | LOC_Os01g66100 | enzyme | plant stature, dwarfism | development | 11961544 |
| OsC1 | LOC_Os06g10350 | TF (R2R3-MYB) | plant coloration (loss of apiculus color) | other | 15514070, 23034647, 20972439 |
| Rc | LOC_Os07g11020 | TF (bHLH) | seed color | other | 17696613, |

| | | | | | |
|---|---|---|---|---|---|
| | | | | | 16399804, 20972439 |
| Waxy /GBSS | LOC_Os06g04200 | enzyme | amylose content (glutinous rice) | other | 7742858, 9718725, 9747848, 16547098 |
| Bh4 | LOC_Os04g38660 | transporter | seed hull color | other | 21263038, 23034647 |
| Pikm1-TS + Pikm2-TS cluster | LOC_Os11g46210 | R-protein with leucine-rich repeats | pathogen resistance | biotic stress response | 18940787, 22643901 |
| Pikm1-TS + Pikm2-TS cluster | LOC_Os11g46200 | R-protein with leucine-rich repeats | pathogen resistance | biotic stress response | 18940787, 22643901 |
| Pi5-1 + Pi5-2 cluster | LOC_Os09g15840 | R-protein with leucine-rich repeats | pathogen resistance | biotic stress response | 19153255 |
| Pib | LOC_Os02g57310 | R-protein with leucine-rich repeats | pathogen resistance | biotic stress response | 10417726 |
| Hd1 | LOC_Os06g16370 | TF CO-like | flowering time | development | 11148291, 19246394,15078816, 19246394 |
| SaM + SaF | LOC_Os01g39670 | E3 Ligase + F-box | F1 male sterility | other | 19033192 |
| SaM + SaF | LOC_Os01g39680 | E3 Ligase + F-box | F1 male sterility | other | 19033192 |

**Supplementary Table S3** Causal gene features and their frequency and value in causal genes and the genome background

| Category | Feature name[a] | Arabidopsis | | | Rice | | |
|---|---|---|---|---|---|---|---|
| | | causal gene | genome gene | P value[b] | causal gene | genome gene | P value[b] |
| Enzyme class | is_carbohydrates_metabolism | 1.7% | 5.0% | 3.7E-01 | 6.7% | 5.2% | 5.1E-01 |
| Enzyme class | is_nucleotides_metabolism | 0.0% | 2.2% | 6.4E-01 | 2.2% | 2.5% | 1.0E+00 |
| Enzyme class | is_energy_metabolism | 3.3% | 1.9% | 3.1E-01 | 0.0% | 1.8% | 1.0E+00 |
| Enzyme class | is_fatty_acids_lipids_metabolism | 3.3% | 2.8% | 6.9E-01 | 0.0% | 3.2% | 4.0E-01 |
| Enzyme class | is_specialized_metabolism | 5.0% | 3.7% | 4.8E-01 | 0.0% | 4.4% | 2.7E-01 |
| Enzyme class | is_cofactors_metabolism | 0.0% | 2.1% | 6.4E-01 | 2.2% | 2.4% | 1.0E+00 |
| Enzyme class | is_other_metabolism | 0.0% | 1.3% | 1.0E+00 | 2.2% | 1.3% | 4.4E-01 |
| Enzyme class | is_hormones_metabolism | 1.7% | 1.2% | 5.2E-01 | 6.7% | 1.2% | 1.8E-02 |
| Enzyme class | is_inorganic_nutrients_metabolism | 8.3% | 1.4% | 1.7E-03 | 2.2% | 1.9% | 5.9E-01 |
| Enzyme class | is_detoxification_metabolism | 0.0% | 0.9% | 1.0E+00 | 0.0% | 1.1% | 1.0E+00 |
| Enzyme class | is_redox_metabolism | 0.0% | 0.7% | 1.0E+00 | 0.0% | 1.0% | 1.0E+00 |
| Enzyme class | is_amino_acids_metabolism | 1.7% | 1.7% | 1.0E+00 | 0.0% | 1.9% | 1.0E+00 |
| Enzyme class | is_macromolecule_metabolism[c] | 11.7% | 12.9% | 1.0E+00 | 44.4% | 33.9% | 1.6E-01 |
| GO | is_TF | 10.0% | 6.3% | 2.8E-01 | 13.3% | 4.8% | 2.0E-02 |
| GO | is_receptor | 11.7% | 0.6% | 1.5E-07 | 0.0% | 0.6% | 1.0E+00 |
| GO | is_kinase | 15.0% | 10.9% | 3.0E-01 | 17.8% | 13.3% | 3.8E-01 |

| | | | | | | | |
|---|---|---|---|---|---|---|---|
| GO | is_transporter | 20.0% | 7.9% | 2.4E-03 | 11.1% | 7.1% | 2.5E-01 |
| Co-function network | network_weight [d] | 182.0 | 126.4 | 5.9E-02 | 11.8 | 76.5 | 1.2E-21 |
| Polymorphism | is_start_gained | 56.7% | 43.0% | 3.7E-02 | 11.1% | 11.8% | 1.0E+00 |
| Polymorphism | is_start_lost | 6.7% | 7.3% | 1.0E+00 | 0.0% | 0.8% | 1.0E+00 |
| Polymorphism | is_stop_gained | 68.3% | 35.8% | 4.6E-07 | 11.1% | 5.4% | 9.4E-02 |
| Polymorphism | is_stop_lost | 11.7% | 8.0% | 3.3E-01 | 4.4% | 1.3% | 1.2E-01 |
| Polymorphism | is_SNP_splice_site | 28.3% | 19.5% | 1.0E-01 | 24.3% | 23.5% | 8.5E-01 |
| Polymorphism | Is_SNP_cis [e] | 55.0% | 72.0% | 5.7E-03 | NA | NA | NA |
| Polymorphism | is_nonsyn_deleterious [f] | 93.3% | 82.5% | 2.6E-02 | 40.0% | 28.7% | 1.0E-01 |
| Polymorphism | normalized_nonsyn_SNP [g] | 0.159 | 0.113 | 9.1E-03 | 0.011 | 0.011 | 4.7E-01 |
| Evolution related | is_essential_gene [h] | 1.7% | 10.0% | 2.8E-02 | 0.0% | 8.9% | 3.1E-02 |
| Evolution related | paralog_copy_number | 56.3 | 30.4 | 9.9E-07 | 118.0 | 37.0 | 1.5E-03 |

[a] The features with a name that starts with "is_" are binary variables.
[b] Fisher's exact test was used for all binary features. Mann-Whitney U Test was used for all continuous variables.
[c] This category contains genes that were annotated as enzyme by GO but not present in PMN. A majority of them are involved in macromolecule metabolism.
[d] "network_weight" was the sum weight of all edges connected to a gene based on AraNet or RiceNet (Lee *et al.* 2010; Lee *et al.* 2011).
[e] "is_SNP_cis" indicates whether polymorphisms are present in the cis-element found in the promoter region of the gene (Weirauch *et al.* 2014). The cis-element data was only available for Arabidopsis and therefore not used in the rice model.
[f] "is_nonsyn_deleterious" indicates the presence of non-synonymous SNPs in highly conserved residues, which are likely to affect protein function (Ng and Henikoff 2003).
[g] The number of non-synonymous SNPs was normalized to the length of the protein.
[h] Essential gene prediction was based on function annotation, duplication, expression levels and patterns, rate of evolution, cross-species conservation and other information and taken from (Lloyd *et al.* 2015).

**Supplementary Table S4** Features used for the Arabidopsis model. Known causal genes were labeled as 1 in the 'class' column. Other genes were labeled as 0.

This file is available as 'Arabidopsis_features_v4.csv' at

https://github.com/carnegie/QTG_Finder/tree/master/prediction/model_training

**Supplementary Table S5** Features used for the rice model. Known causal genes were labeled as 1 in the 'class' column. Other genes were labeled as 0.

This file is available as 'rice_features_v2.csv' at
https://github.com/carnegie/QTG_Finder/tree/master/prediction/model_training

**Supplementary Table S6** Evaluating model performance based on the rank of an independent set of known causal genes.

| QTL trait | Gene name | Rank in a QTL[a] | Evidence code [b] | Reference |
|---|---|---|---|---|
| **Arabidopsis** | | | | |
| Seed size | *AHK3* | 1/366 (<0.1%) | G | Guo *et al.* ,2016<br>Riefler *et al.*, 2006 |
| Seed size | *AHK2* | 2/293 (<0.1%) | G | Guo *et al.* ,2016<br>Riefler *et al.*, 2006 |
| Stem branching | *MAX3* | 17/457 (3%) | G, J | Huang *et al.*, 2013<br>Ehrenreich *et al.*, 2007<br>Bennett *et al.*, 2006 |
| Insect resistance | *TGG2* | 31/496 (6%) | G, J | Pfalz *et al.*, 2007<br>Barth *et al.*, 2006 |
| Insect resistance | *TGG1* | 38/496 (7%) | G, J | Pfalz *et al.*, 2007 |

| | | | | |
|---|---|---|---|---|
| | | | | Barth *et al.*, 2006 |
| Insect resistance | *GSOH1* | 15/104 (13%) | C, E, G | Hansen *et al.*, 2008 |
| Seed germination | *FBA2* | 23/111 (19%) | G | Yuan *et al.*, 2016 |
| Stem branching | *AGL6* | 106/457 (22%) | C, G, H | Huang *et al.*, 2012 |
| Fructose content in leaves | *Sweet17* | 47/186 (24%) | C, F, G | Fabien *et al.*, 2013 |
| Seed germination | *AZF2* | 38/111 (33%) | G | Yuan *et al.*, 2016 |
| Shoot regeneration | *RPK1* | 282/554 (50%) | H, G, J | Motte *et al.*, 2014 |
| **Rice** | | | | |
| Blast resistance | Pi35/Pish | 3/203 (0.5%) | C, G, H | Fukuoka *et al.*, 2014 |
| Blast resistance | Pi-k | 10/444 (2%) | E, C, G | Sharma *et al.*, 2005<br>Azizi *et al.*, 2016<br>Rai *et al.*, 2011 |
| Leaf blade width | nal1 | 7/210 (2%) | C, G | Qi *et al.*, 2008 |
| Grain chalkiness | *qACE9* | 9/120 (6%) | F, E | Gao *et al.*, 2016 |
| Plant height | osh15 | 14/117 (11%) | F, G | Fan *et al.*, 2016<br>Sato *et al.*, 1999 |
| Heading time | Hd3a | 102/730 (13%) | F, G | Monne *et al.*, 2002<br>Itoh *et al.*, 2010 |
| Grain color | kala4 | 15/62 (22%) | F, G | Oikawa *et al.*, 2015 |
| Tiller number | ts1 | 54/175 (30%) | F, E | Liu *et al.*, 2017 |
| Bacterial blight resistance | *WRKY13* | 171/460 (36%) | G | Hu *et al.*, 2008<br>Qiu *et al.*, 2007 |
| Blast and bacterial blight lesion | UROD | 19/38 (47%) | F | Zeng *et al.*, 2013 |

[a] The numerator is the rank of the causal gene. The denominator is the total number of genes in the QTL region. The percentage in parentheses indicates rank percentile.

[b] Evidence code: C, functional complementation; E, association between the allelic status and expression of the gene; F, fine mapping; G, genetic analyses; H, haplotype analyses showing the association between the trait and polymorphisms of the gene. J, joint linkage-association analyses

#### Supplementary Figures

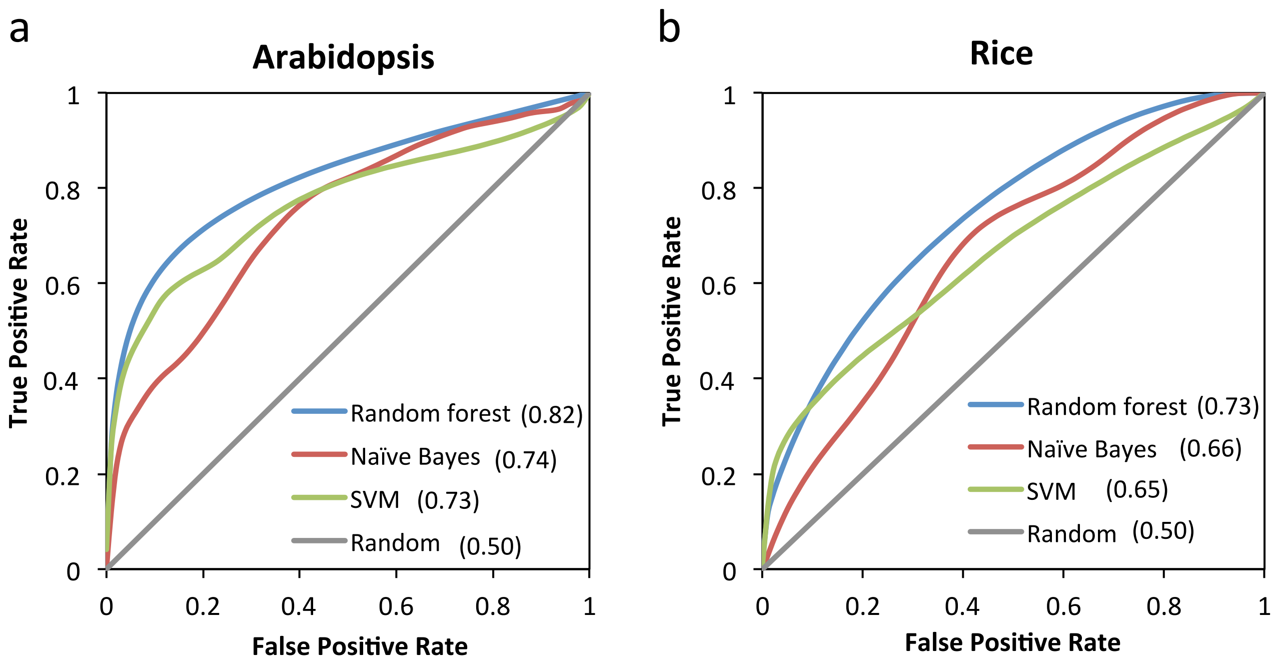


**Supplementary Fig. S1** Comparing ROC curves of classifiers. ROC curves were based on cross-validation. Numbers in parentheses indicate Area Under the Curve (AUC). Grey diagonal lines indicate the expected training performance of a model based on random guessing. (a) Arabidopsis (b) rice

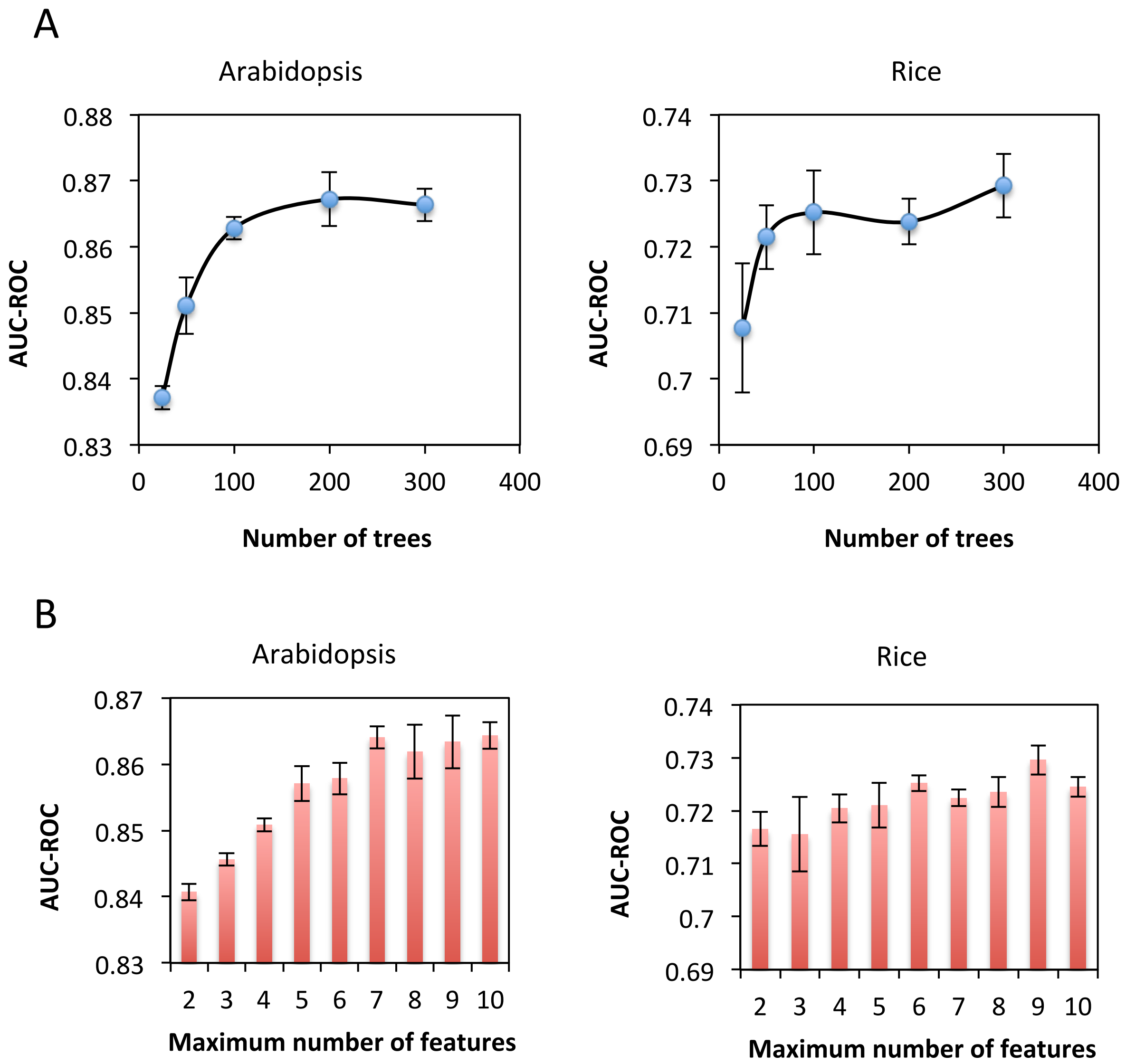


**Supplementary Fig. S2** Random forest parameters. Area Under the Curve of ROC (AUC-ROC) were based on cross-validation. Error bars indicate standard deviation. (a) Number of trees in the forest (b) The maximum number of features to consider when looking for the best split.

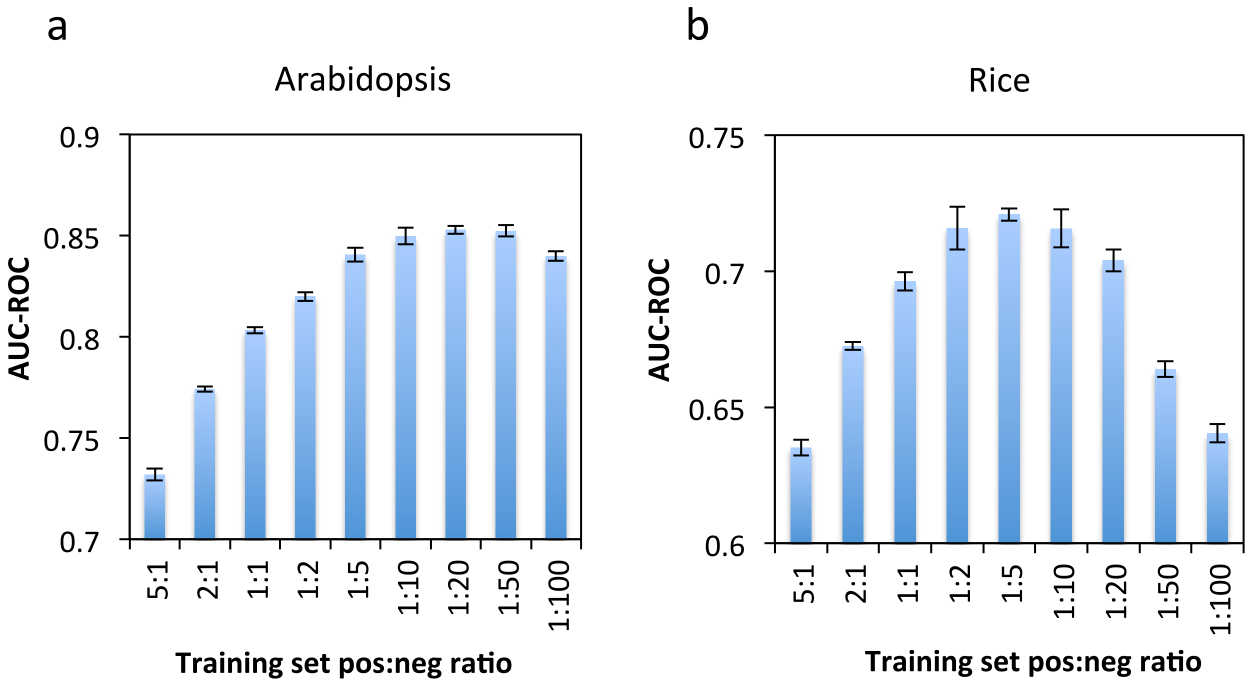


**Supplementary Fig. S3** The ratio of positives:negatives in the training set. Area Under the Curve of ROC (AUC-ROC) were based on cross-validation. Error bars indicate standard deviation. (a) Arabidopsis (b) rice

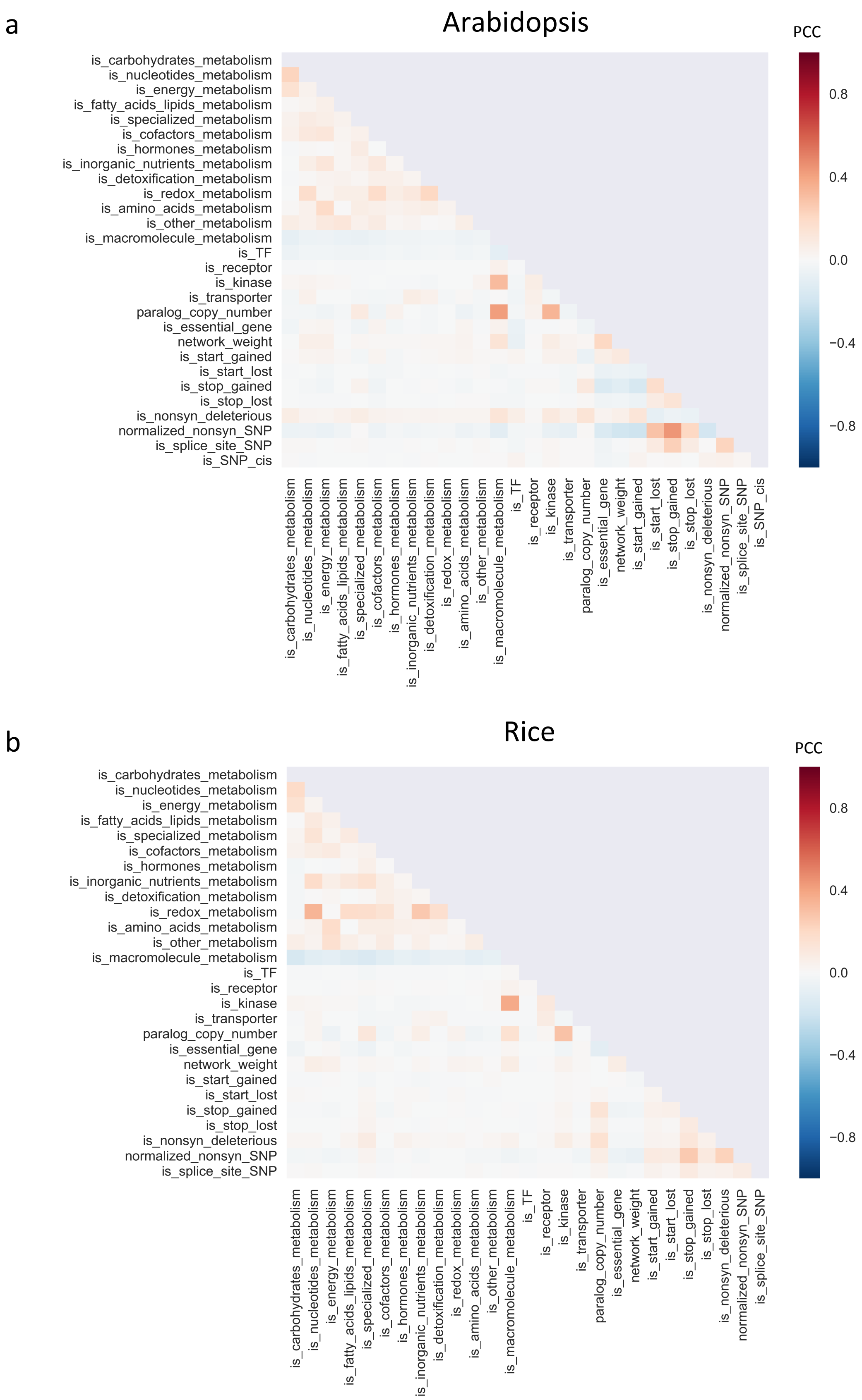


**Supplementary Fig. S4** Correlations among features. Pearson's correlation coefficients (PCC) were used for the heatmap. (a) Arabidopsis (b) rice

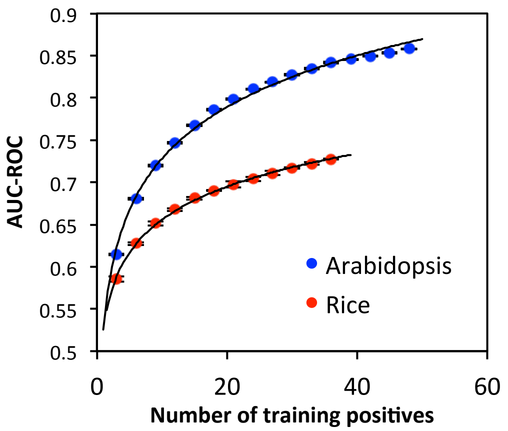


**Supplementary Fig. S5** The relationship between training performance and the number of training positives used. The cross-validation framework was applied to training sets with different number of training positives. The number of training negatives was adjusted to maintain the same positive:negative ratio in all training sets. One fifth of known causal genes were retained as testing positive to calculate AUC-ROC.
